# Meiotic double strand DNA breaks and spontaneous mutation in *Drosophila melanogaster*

**DOI:** 10.1101/2025.10.29.685400

**Authors:** Rob Melde, Austin Daigle, JoHanna Abraham, Nathaniel Sharp

**Affiliations:** Department of Genetics, University of Wisconsin Madison; Department of Genetics, University of North Carolina at Chapel Hill

## Abstract

The exchange of genetic material during meiosis requires the formation and repair of DNA double-strand breaks (DSBs), which may not be repaired with perfect fidelity. If meiotic exchange is mutagenic, this would add to the costs of sexual reproduction and affect patterns of genome evolution, but much of the evidence for this is indirect. In the fruit fly *Drosophila melanogaster*, it is possible to completely suppress endogenous DSBs while retaining normal fertility. We took advantage of this system to generate fly strains with and without a mutant allele of *mei-P22*, a gene that is essential for meiotic DSB formation, on a common genetic background. This allowed us to investigate the relationship between DSBs and genome-wide mutation patterns, using a mutation accumulation design to allow un-selected spontaneous mutations to be observed. Following 30 generations of mutation accumulation, we identified over 1800 mutations by whole-genome sequencing. The presence of meiotic DSBs had little effect on the rate and spectrum of point mutations. We found that mutations were more likely to occur in areas of the genome with higher rates of crossover recombination, regardless of whether meiotic DSBs were occurring. We also found that the rate of transposable element insertions across multiple TE families was substantially elevated in the group lacking meiotic DSBs, suggesting that host suppression of mobile genetic elements is closely associated with meiotic recombination mechanisms.

## Introduction

Meiotic recombination (MR) is a key driving force of evolution. MR is ubiquitous across eukaryotes and is a core aspect of sexual reproduction. While MR is generally considered evolutionarily favorable (Keightley & Otto, 2006), these benefits come with costly tradeoffs. One obvious cost of MR is the time and energy cells must spend to perform this process. For example, in the model yeast *Saccharomyces cerevisiae*, the mitotic cell cycle takes approximately 90 minutes under optimal conditions, whereas meiosis and sporulation require at least 8-12 hours to complete (Duina et al., 2014; Ray & Ye, 2013). From a population genetics perspective, beneficial alleles that work well in concert with one another can be broken apart by MR, slowing adaptation (Charlesworth & Barton, 1996; Charlesworth & Charlesworth, 1975). At the molecular level, while MR typically leads to proper chromosome segregation, errors in this process can lead to chromosomal nondisjunction and aneuploidy (Koehler et al., 1996). Another potential molecular cost relates to the formation and repair of meiotic double strand DNA breaks (DSBs), which are required for MR. Some DSBs occur spontaneously as a result of DNA damage; these events, described as “non-programmed” or “exogenous” DSBs, are considered highly mutagenic because error prone pathways are commonly used to repair them (Rodgers & McVey, 2016; Strathern et al., 1995). In contrast, the “endogenous” or “programmed” DSBs that occur during meiosis are thought be more tightly regulated and repaired using specialized pathways (Longhese et al., 2009; Wei et al., 2019). Nevertheless, endogenous DSBs likely contribute at least some additional risk of mutation, and there is some evidence for such an effect.

A positive correlation between crossover recombination rate and molecular diversity may be an indirect source of evidence on the relationship between endogenous DSBs and mutation, but this pattern is not observed in all species, and can instead be explained by reduced background selection in areas of higher crossover recombination (Begun & Aquadro, 1992; Cutter & Payseur, 2013; Smukowski & Noor, 2011). Stronger evidence of a relationship between MR and mutation comes from humans, where SNPs and *de novo* germline mutations are found more commonly near known recombination hotspots (Hinch et al., 2023; Pratto et al., 2014). Similarly, short insertions and deletions (INDELs) and structural variants have been shown to be associated with recombination hotspots in human pedigree studies (Beyter et al., 2021; Hinch et al., 2023). Parent-offspring sequencing has also revealed an association between crossover events and germline mutations in humans (Halldorsson et al., 2019) and to a lesser extent in bees (Liu et al., 2017; Yang et al., 2015). Parent-offspring sequence data in *Drosophila melanogaster* and *D. simulans* did not detect any relationship between crossovers and germline mutations, but the number of relevant phased mutations was low (14 female-derived mutations) (Wang et al., 2023). In *S. cerevisiae*, meiosis is associated with a higher mutation rate than mitosis (Magni, 1963), and this effect has been directly linked to endogenous DSBs (Rattray et al., 2015). Additional direct tests of the impact of MR on the rate and spectrum of point mutations would be valuable.

Another mutational process associated with recombination is the insertion of transposable element (TE) sequences. TEs are found more often in genomic regions of low recombination (Kent et al., 2017; Kofler et al., 2012; Petrov et al., 2011). The Hill-Robertson effect and selection against ectopic recombination could drive this pattern (Biemont et al., 1997; Dolgin & Charlesworth, 2008; Kelleher et al., 2020), but there is also evidence that TEs suppress recombination (Huang et al., 2025). The formation of endogenous DSBs has been shown to be related to both TE mobility and TE silencing (Dufourt et al., 2014; Wei et al., 2019; Wylie et al., 2016). This suggests that altering the formation of endogenous DSBs could influence TE mobility in an activating or suppressing manner, but this possibility has not been thoroughly explored.

Quantifying the mutational consequences of endogenous DSBs is challenging. In many organisms, endogenous DSBs are essential for proper chromosomal segregation and gamete formation (Petronczki et al., 2003). In such systems, researchers are typically limited to comparing mutation between mitotically and meiotically dividing cells or examining mutational patterns near crossover events and recombination hotspots, which are approaches with significant limitations. Comparing mitotic versus meiotic cells confounds the effects of DSBs with the broader differences between fundamentally distinct cell division processes. Additionally, focusing on crossover-associated mutations provides only a partial view of mutations that may be associated with endogenous DSBs, since many meiotic DSBs are resolved without crossing over. These methodological constraints therefore limit our understanding of how endogenous DSBs truly impact the overall mutational landscape.

Conveniently, *D. melanogaster* can perform meiosis and form viable gametes without endogenous DSBs. In nearly all *Drosophila* species, males naturally perform meiosis without endogenous DSBs or crossover recombination (Lee & Rosin, 2024). Laboratory strains have been developed where meiotic DSBs are also absent in the female germline. One example contains the allele *mei-P22[P22]* (also known as *mei-P22[1], mei-P22[McKim]* and henceforth referred to as *mei-P22^−^*), generated with P-element mutagenesis (Sekelsky et al., 1999). The *mei-P22* protein forms a dimer with *mei-W68* (a paralog of *spo-11*) and interacts with *Trem*, *Vilya,* and *Narya* to form endogenous DSBs (Lake et al., 2011, 2015, 2019; Liu et al., 2002; Mehrotra & McKim, 2006). The *mei-P22^−^* allele effectively eliminates transcription of *mei-P22*, preventing endogenous DSB formation and MR in the female germline while permitting viable gamete production (Iida & Lilly, 2004; Liu et al., 2002; McKim et al., 1998). Importantly, the effect of *mei-P22^−^* is distinct from that of balancer chromosomes, which inhibit crossing over but do not prevent DSBs (Miller et al., 2016).

In *D. melanogaster* females, meiosis involves about six crossovers throughout the genome, but a total of 20-24 endogenous DSBs that must be repaired (Carpenter, 1987; Mehrotra & McKim, 2006). Compared to organisms with recombination hotspots, the landscape of crossing over in *D. melanogaster* females is relatively uniform. The correlation between mutation and recombination found in humans may be more apparent than such a relationship in flies would be, because there are specific 1-2 kb regions where recombination (and thus endogenous DSBs) tend to occur, mediated by PRDM9 hotspots (Baudat et al., 2010; Jeffreys et al., 2001). Endogenous DSBs in *D. melanogaster* have only been observed to occur within euchromatin, and are influenced by accessible chromatin states linked to active transcription but are otherwise broadly distributed throughout the genome (Adrian et al., 2016; Adrian & Comeron, 2013; Mehrotra & McKim, 2006).

To study the effect of meiotic DSBs on mutation, we established two sets of mutation accumulation (MA) lines: one using females undergoing normal endogenous double-strand break (DSB) formation, and another using *mei-P22*^−^ females in which endogenous DSBs were absent. In MA experiments, repeated bottlenecking minimizes the efficacy of natural selection, allowing most new mutations to become fixed with neutral probability (reviewed in Katju & Bergthorsson, 2019). Such experiments are a powerful means of generating large, unbiased mutation datasets with relatively modest sequencing effort. We used this approach to compare the pattern of germline mutation in the presence and absence of endogenous meiotic DSBs. We found that eliminating endogenous DSBs had little impact on the rate and spectrum of point mutations but caused a significant increase in the rate of transposable element insertion throughout the genome.

## Methods

### Fly stocks and mutation accumulation

We reared flies on defined medium (14.3 g/L agar, 92.3 g/L white sugar, 46 g/L debittered yeast, 7.4 g/L potassium sodium tartrate, 0.93 g/L potassium phosphate, 0.46 g/L sodium chloride, 0.46 g/L calcium chloride, 0.46 g/L magnesium chloride, 0.46 g/L iron sulfate, 0.5% propionic acid) in standard vials seeded with live yeast, incubating at 25 C with a 12-hour light:dark cycle. We obtained stock 4931 from the Bloomington Drosophila Stock Center (RRID:BDSC_4931) which is *y*^1^ *w*^1^/Dp(1;Y)*y*^+^; P{*w*[+mC]=lacW}*mei-P22^P22^*; *sv^spa-pol^*. To ensure a fair comparison of the mutation patterns between treatment groups we sought to establish MA lines with and without *mei-P22^−^* from the same genetic background. To create these lines we first backcrossed *mei-P22^−^* into a stock of *w*^1^ *y*^1^/Dp(1;Y)*y*^+^ for five generations; *mei-P22^−^* can be tracked in these crosses because it is marked with a mini-white allele, which partially reverses the eye color effect of *w*^1^ (*mei-P22^−^* does not eliminate crossing over in the heterozygous state). In this genetic background, XX females are *y*^1^/*y*^1^ and express a yellow body color phenotype; XY males are *y*^1^/Dp(1;Y)*y*^+^, where Dp(1;Y)*y*^+^ marks the Y chromosome with a functional copy of *y*, such that males express a wild type body color. The presence of *mei-P22^−^* increases the rate of non-disjunction (Liu et al., 2002; Sekelsky et al., 1999), resulting in viable XXY females and sterile X∅∥ males, which can be identified by body color (wild type body color in XXY females and yellow body color in X∅∥ males). After backcrossing for five generations, we imposed single-pair bottlenecks for an additional five generations to remove standing genetic variation. We then performed a final set of expansion crosses, selecting either for or against *mei-P22*^−^, establishing 20 lines in each treatment group. For MA, each line was bottlenecked for 30 consecutive generations using one randomly selected male and one randomly selected female, excluding XXY and X∅∥ flies. Five lines went extinct during the course of the experiment, which ultimately concluded with 16 lines where endogenous DSBs occurred (*mei-P22*^+^) and 19 lines where they did not (*mei-P22*^−^).

### DNA Extraction and Sequencing

Upon completion of the 30^th^ generation of MA, we collected all female offspring (6-24 individuals per line) and froze them at –80 C. We extracted genomic DNA from stored flies using the Qiagen DNeasy Blood & Tissue Kit with the insect protocol (catalog #69504), purified using the Qiagen DNeasy PowerClean Kit (catalog #12877-50), and quantified using a Qubit Fluorometer (ThermoFisher Scientific). We submitted DNA to the University of Wisconsin Biotechnology Center DNA Sequencing Core Facility (RRID:SCR_017759) for paired-end whole genome sequencing using an Illumina NovaSeq X Plus.

### Calling point mutations

We obtained about 2.3 billion reads in total, which we quality checked using FastQC (v0.12.1) and trimmed using Trimmomatic (v0.39) (Andrews, 2010; Bolger et al., 2014). After masking repetitive regions using RepeatMasker (v4.1.5), we mapped reads to the *D. melanogaster* reference genome (v6.59) using bwa-mem2 (v2.2.1) (Dos Santos et al., 2015; Smit et al., 2013; Vasimuddin et al., 2019), resulting in a mean depth of coverage of 99. We sorted and indexed BAM files using Samtools (v1.6), removed duplicate reads using Picard Tools MarkDuplicates (v3.1.1), and called variants using GATK HaplotypeCaller (v4.5.0.0) (Broad Institute, 2019; Li et al., 2009). Finally, we used Rstudio (v4.3.2) for variant filtering, statistical modeling, and figure creation (R Core Team, 2023; Van der Auwera & O’Connor, 2020). We utilized glmmTMB (v1.1.11; Brooks et al., 2017) to fit generalized linear mixed effect models of genome wide mutation patterns.

To filter our callset we first selected variants that were only present in one line and required that the site of interest have a called genotype in all lines (finding the exact same mutation in two or more lines due to convergent mutation is highly unlikely, and such cases should represent sequencing errors or preexisting variation). We required there to be at least 10 supporting reads for the variant allele and considered only variants passing the filters recommended by GATK: mapping quality ≥ 50, quality by depth ≥ 2, Fisher strand bias ≤ 60, strand odds ratio ≤ 3, absolute mapping quality rank sum ≤ 8, and absolute read position rank sum ≤ 4. We additionally required that depth at a variant site was no less than half and no greater than twice the median chromosome-wide depth in that line. Finally, we removed any single-nucleotide variants (SNV) or INDELs called within 1 kb of a structural variant (SV) or TE insertion. Where possible, we applied the same filtering criteria to non-variant sites when calculating the number of callable sites in the genome. For a site to be considered callable we required that site to pass the filters in all lines. We considered variants only on the major contigs listed in the *D. melanogaster* reference genome that are present in female flies and that normally recombine (2L, 2R, 3L, 3R, and X; variants called on chromosome 4 are also listed in Supplementary Material). The reference genome for these contigs includes 133,880,608 sites and our final callset includes 99,458,130 sites (74%). Of the 34,422,478 non-callable sites, 20,052,895 were not considered callable because they were deemed repetitive by RepeatMasker. The remaining 14,369,583 sites were considered non-callable because they failed one or more of our filtering metrics. We categorized events as single nucleotide variants (SNV), short insertions and deletions (INDEL), multi-nucleotide variants (MNV; two or more SNV within 1 kb in the same line), and complex variants (COMPLEX; two or more variants within 1 kb in the same line where at least one variant was an INDEL). For the purposes of mutation rate estimation, each MNV and COMPLEX group of variants was counted only once. In order to understand the predicted functional impacts of the variants in our dataset, we used Ensemble’s variant effect predictor (release 113) to identify the most severe effect predicted for each variant (McLaren et al., 2016). We set the upstream/downstream distance to zero and kept all other parameters at their defaults.

We estimated mutation rates using both homozygous and heterozygous variants, and dividing by 30 MA generations. By including heterozygous mutations, this procedure will overestimate the true mutation rate, since more than one copy of a given chromosome is effectively included in each line. We did not find evidence that the proportion of mutations that are heterozygous differs between treatment groups (Results), and so we don’t expect this bias to affect our treatment comparison. Considering only homozygous mutations would produce the opposite bias, since mutations cannot become homozygous immediately. Performing our statistical comparisons using only homozygous mutations did not alter our conclusions (supplemental R script).

### Calling structural variants

To identify larger structural variants that may have been missed by HaplotypeCaller we used MANTA (v.1.6.0) (X. Chen et al., 2016). For practical purposes we analyzed our 35 samples in 3 blocks (10, 10 and 15 samples) and then combined the outputs. Each analysis block had lines from both treatment groups. We then filtered variants by selecting only those that were present in one line with a called genotype for every line present in that block. We removed any variants that did not have at least 10 supporting spanning reads or 10 supporting split reads, and considered only cases with the maximum possible quality score.

### Calling transposable element variants

We used the McClintock meta-pipeline (v.2.0.0) to detect transposable element (TE) insertions and excisions using multiple detection methods (J. Chen et al., 2023; Nelson et al., 2017), specifically, the McClintock implementation of TEMP2 (Yu et al., 2021) employing the default settings from McClintock. Because TE callers rely on information from discordant read pairs, where one read in the pair maps to the reference genome and the other maps to a TE sequence, several studies have shown that trimming reads can improve the detection accuracy of these tools, despite a reduction in coverage depth (Daigle et al., 2025; Yu et al., 2021). This is especially relevant for our dataset, where the median insert size (about 300 bp) is close to the total length of read pairs (150 × 2 bp), which will lead to many overlapping pairs. For this reason, we trimmed all reads at the 3’ end down to 50 bp using fastp (J. Chen et al., 2023) before TE detection. We used the consensus TE sequences and reference genome annotation created by Rech et al. (available at http://doi.org/10.20350/DIGITALCSIC/13765 and http://doi.org/10.20350/DIGITALCSIC/13894, respectively) as inputs to McClintock (Rech et al., 2022).

To identify non-reference TE insertions unique to each MA line, we compared the sets of predicted insertions across all samples. Because TE callers often report slightly different breakpoint coordinates for the same insertion (J. Chen et al., 2023; Daigle et al., 2025), we treated insertions in different lines from the same TE family located within 50 bp of one another as representing the same event. Using this criterion, we identified TE insertions confidently present in only one MA line, which we interpreted as *de novo* mutations specific to that lineage. We excluded non-reference insertions present in more than one line, as it is unclear if these are the result of detection error, recurrent mutation TE insertions at the same site, or variants that remained segregating in the stocks used to create the MA lines. TEMP2 additionally checks for the presence or absence of reference TE sequences. We interpreted reference TEs uniquely absent in one MA line as de novo TE excision events. However, we note that we do not have the ability to distinguish between a deletion that removes a TE and a true enzymatic excision of a TE.

### Analysis of genome wide mutation patterns

We used statistical models to test whether specific genomic features could account for variation in the locations of mutations throughout the genome, and whether these patterns differed between the two treatment groups. We considered crossover recombination rates, replication timing, chromatin state information and G/C content. We used 7657 genomic windows defined by chromatin state (Filion et al., 2010), and added information on crossover recombination rates (Comeron et al., 2012) and replication timing (Schwaiger et al., 2009) for each window, using weighted averaging to account for the different window sizes for each variable. We used the Kc cell type replication timing data in our analyses (Schwaiger et al., 2009), but note that the Kc and Cl8 cell type replication timing values are strongly correlated (*P* < 2.2 × 10^−16^; *R^2^* = 0.812). For each window we used a custom bash script to count the number of sites that were considered callable in our dataset as well as the total number of G/C sites in that window according to the reference genome. The chromatin state data includes five categories (Filion et al., 2010), with two being distinct types of euchromatin, referred to as “red” (RE) and “yellow” (YE), two being distinct types of heterochromatin, referred to as “blue” (BL) and “green” (GR), and one being a highly repressed type of chromatin, referred to as “black” (BK). The replication timing and chromatin state datasets were built under an older release of the *D. melanogaster* reference genome, so we used the flybase Drosophila Sequence Coordinates Converter (https://flybase.org/convert/coordinates) to convert coordinates to reference version r6. We excluded chromosomes Y and 4, any windows missing replication timing or recombination data, as well as any windows with zero callable sites.

This windowed approach allowed us to run generalized mixed effect models with the number of variants observed in that window as the response variable and genomic features as predictors. We scaled recombination rate and G/C content to their respective means to facilitate model convergence. We also included power (the number of callable sites multiplied by the number of lines, scaled linearly) as either an additional predictor or an offset term. To compare treatment groups, we duplicated the window data, included treatment as a predictor, and included window identity as a random effect. We found that including chromosome as a random effect did not improve model fit. We used negative binomial generalized linear mixed effect models (GLMMs) with the nbinom2 family (Hardin & Hilbe, 2007) implemented using glmmTMB (Brooks et al., 2017), which accounts for any overdispersion.

## Results

### Point mutation rate

We identified a total of 991 point mutations across 35 MA lines that accumulated mutations for 30 generations (Fig. 1A). The presence of meiotic DSBs did not detectably influence the rate of any point mutation type (SNV: t-test, *P* = 0.92; INDEL: t-test, *P* = 0.95; MNV: Wilcoxon rank sum test, *P* = 0.25; COMPLEX: Wilcoxon rank sum test, *P* = 0.24), or the distribution of point mutations among chromosome arms (χ^2^ = 1.36, *P* = 0.85). The overall point mutation rate did not vary significantly among chromosome arms (χ^2^ = 4.11, *P* = 0.39), or on the X chromosome versus autosomes (binomial test, *P* = 0.45). The majority of the point mutations we identified were in the homozygous state (55.3%), as expected given that MA took place under full-sib mating. We found that that the frequency of homozygous versus heterozygous variants did not differ significantly between groups (Fisher’s exact test, *P* = 0.11; Fig. S1). Regardless of the treatment group, variants on the X chromosome were more likely to be homozygous than those on the autosomes (Fisher’s exact test, *P* = 2.57 × 10^−4^). We found no evidence that the amount of variance among MA lines differed between treatment groups for SNVs, INDELs, MNVs, COMPLEXs, or SVs (Levene tests, all *P* > 0.25).

**Figure 1.**
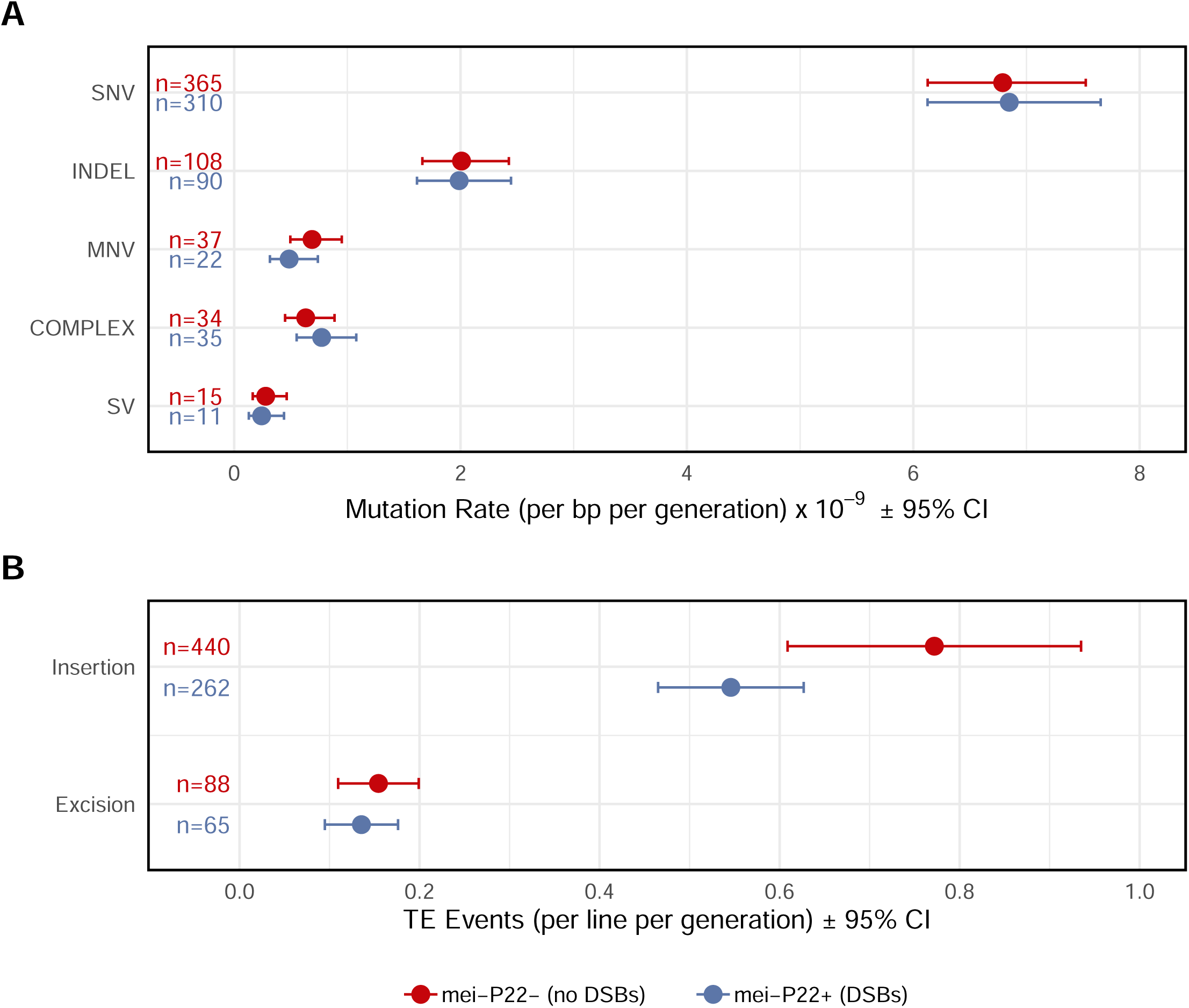
Rates of mutation by group and mutation type. Error bars represent 95% confidence intervals. (a) Rates of single nucleotide variants (SNV), short insertions and deletions (INDEL), multinucleotide variants (MNV), complex variants (COMPLEX), and structural variants (SV) per callable site per generation. (b) Transposable element (TE) insertions and excisions per line per generation.

### Transposable element movement rate

We detected 855 cases of TE movement, of which 82% were insertions. We found a significant difference between treatments in the average number of TE events per line (Welch’s t-test, *t* = 2.35, *P* = 0.027) where the group with endogenous DSBs averaged 0.68 events per line per generation and the group without endogenous DSBs averaged 0.93 TE events per line per generation (Fig. 1B). This result persists when putative TE excisions are excluded (Welch t-test, *t* = 2.43, *P* = 0.022), but not when considering excisions alone (Welch t-test, t = 0.61, *P* = 0.543). We found significantly more variance in the number of TE insertions per MA line in the group that did not perform endogenous DSBs (Levene’s Test, *P* = 7.15 × 10^−3^), but little evidence for a difference in terms of coefficient of variation (bootstrap *P* = 0.083). The group without endogenous DSBs therefore incurred more TE insertion events per generation, with increased among-line variance attributable to the increased mean. In both treatment groups, we observed a significantly higher proportion of TE insertions on the X chromosome than expected based on chromosome size (binomial tests; with endogenous DSBs: *P* = 1.41 × 10^−7^; without endogenous DSBs: *P* = 1.07 × 10^−9^). We detected TEs from numerous different “families” (Fig. 2), including S elements (long inverted repeat DNA transposon), H elements (hAT superfamily DNA transposon), Doc elements (non-LTR LINE retrotransposon) and roo elements (LTR retrotransposon). The increased rate of transposition we observed in the lines lacking endogenous DSBs was evident in most of these TE families (Fig. 2).

**Figure 2.**
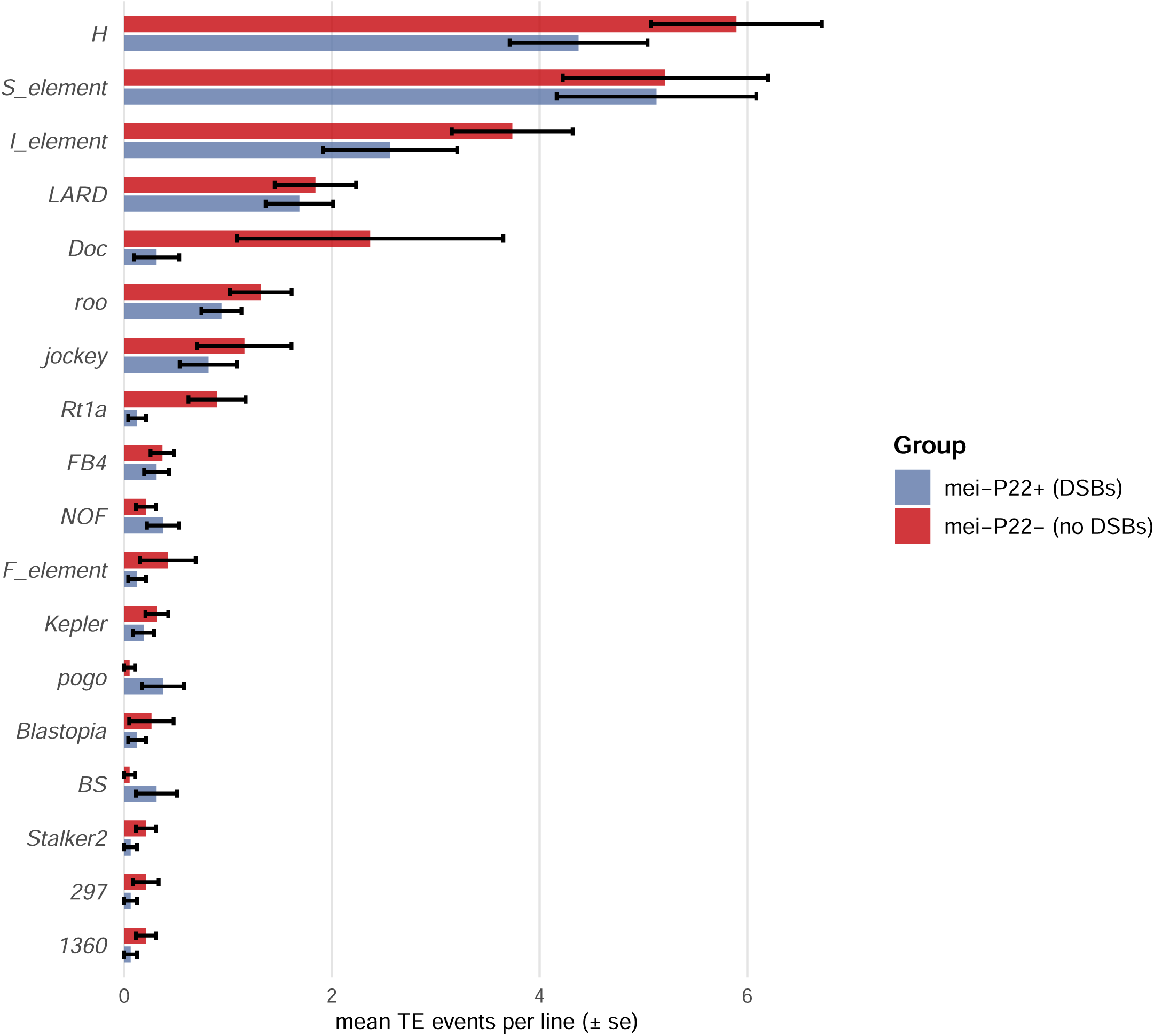
Transposable element activity by family and treatment group. Only TE families with at least five novel insertions or excisions across both treatment groups and at least one insertion/excision in each group are shown here. A complete list is given in Supplementary Material.

### Structural variants

Based on sequencing coverage, we identified one *mei-P22^−^* MA line with apparent trisomy of chromosome 4 (Fig. S2). We also identified 11 large deletions, 13 large insertions, and 2 tandem duplications for a total of 26 structural variants (size 265-7799 bp). The presence of meiotic DSBs did not influence the rate or average length of SVs (Wilcoxon rank sum test, *P* = 0.34; Welch t-test, *t* = 0.11, *P* = 0.91). The distribution of structural variants across chromosome arms did not differ from the null expectation (χ^2^ = 6.52, *P* = 0.16).

### Mutation spectrum

The presence of meiotic DSBs did not significantly influence the SNV spectrum (Fig. 3A; χ^2^ = 6.87, *P* = 0.23) or the transition-transversion ratio (Fig. 3B, Fisher’s exact test, *P* = 0.19). In both treatment groups, SNVs were much more likely to occur at G/C sites than at A/T sites (binomial tests, all *P* < 3.85 × 10^−11^). However, we found some evidence that the strength of this effect differed between treatments (χ^2^ = 4.03, *P* = 0.045), with greater bias towards mutation at G/C sites in the group without meiotic DSBs (69.59% of SNVs versus 62.26%, Fig. 3C). The INDEL mutations in our dataset showed a deletion bias (Fig. 3D; binomial test, *P* = 5.84 × 10^−4^), with no difference in this bias between treatment groups (Fisher’s exact test, *P* = 0.53). The median deletion size was 6 bp and the median insertion size was 15 bp, unaffected by treatment (Wilcoxon rank sum tests, all *P* > 0.30).

**Figure 3.**
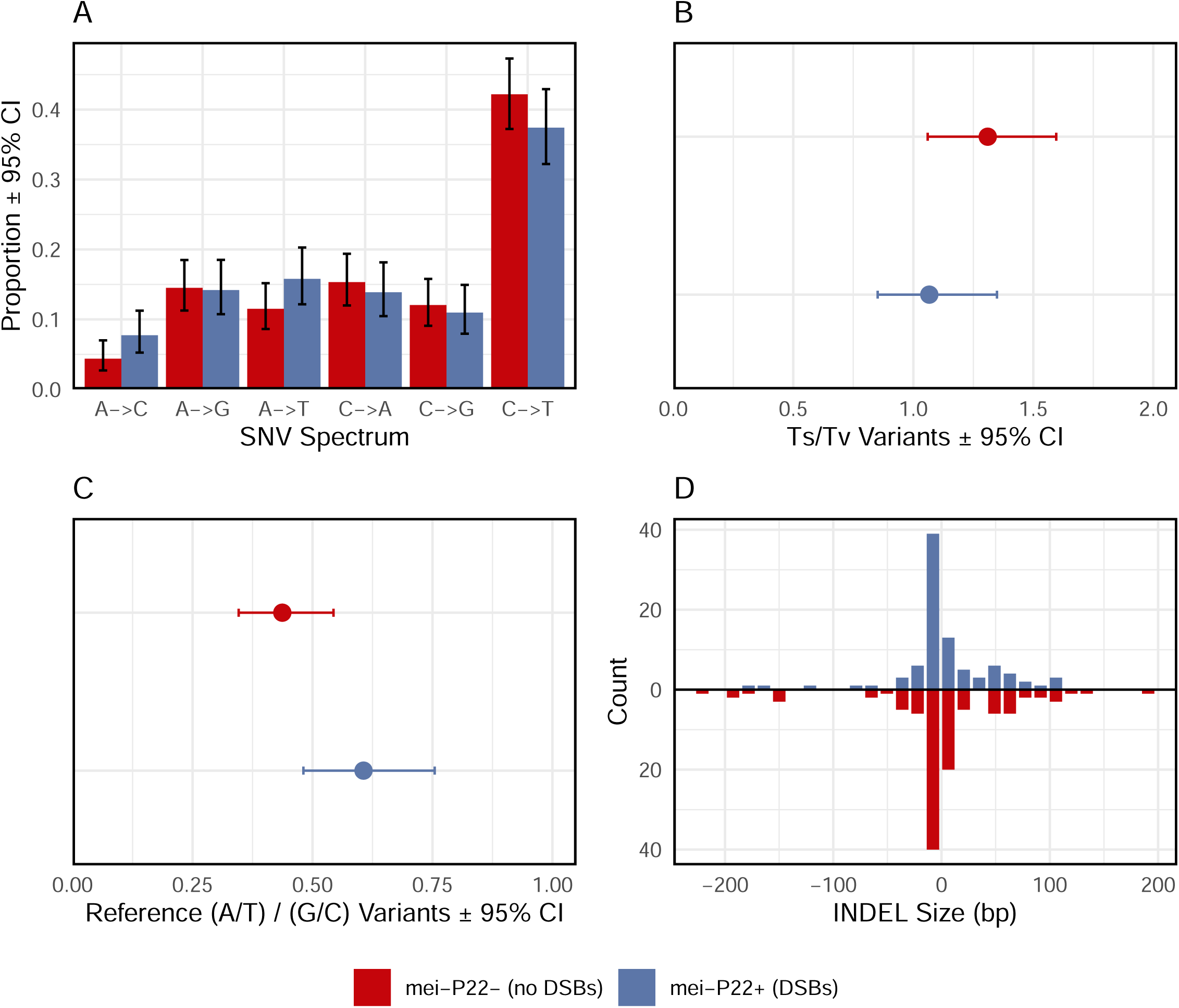
Point mutation spectrum in each treatment group. (a) The spectrum of single nucleotide variants. 95% confidence intervals reflect Wilson score intervals for multinomial proportions. (b) The ratio of transitions to transversions with bootstrap 95% confidence intervals. (c) The ratio of variants that occurred at A/T sites versus G/C sites, with bootstrap 95% confidence intervals. (d) Length distribution of short insertions and deletions.

To ensure that selection did not bias our results, we examined the frequency of mutations in coding regions and their consequences for protein sequences. We found that 72.7% of SNVs and 74.4% of INDELs occurred within genes (including introns; Fig. 4A), which does not significantly differ from a null expectation of 75.08% for our callable sites (binomial tests, *P* = 0.17 and *P* = 0.94, respectively). Similarly, 71.0% of genic SNVs were nonsynonymous, which does not differ significantly from the expected value of 74.4% (binomial test, *P* = 0.38; Fig. 4B). Additionally, these metrics did not differ between treatment groups (Fisher’s exact tests of genic frequency; SNV: *P* = 0.07; INDEL: *P* = 0.87; Fisher’s exact test of nonsynonymous SNVs: *P* = 0.85). We therefore find no evidence that selection biased the accumulation of mutations overall, or affected the treatment groups differently.

**Figure 4.**
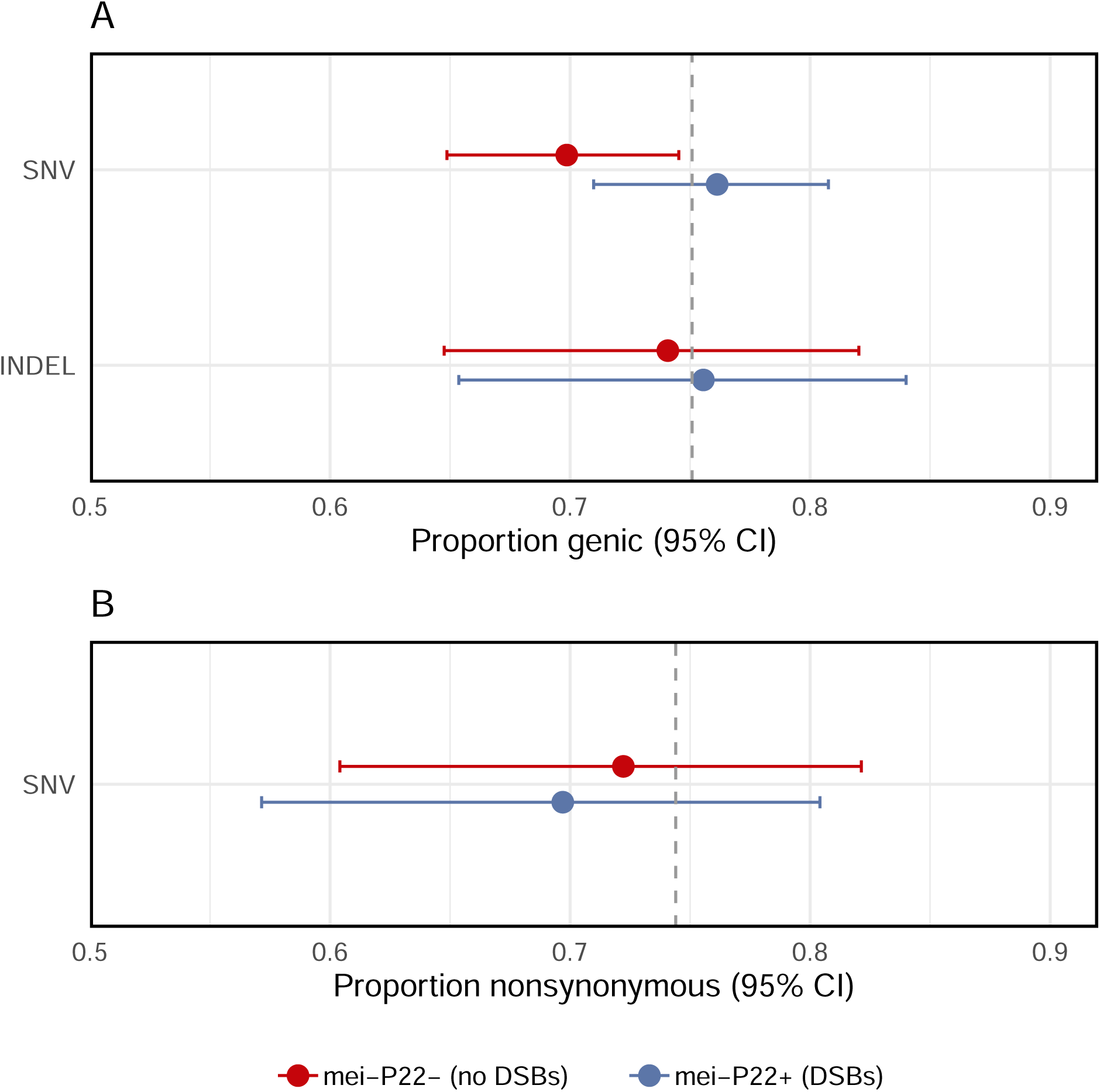
Genomic consequences of point mutations. (a) Proportion of variants found within genes (including introns). The grey dashed line is the proportion of callable sites in our dataset that were genic. (b) Proportion of coding SNVs that were nonsynonymous. The grey dashed line is the null value calculated in Sharp et al. (2016).

### Genomic context of mutations

The presence of endogenous DSBs could affect disproportionately mutation patterns in specific regions of the genome. Considering models of point mutations and TE activity across 7657 genomic windows, we found no evidence that treatment group interacted with G/C content, crossover recombination rate, chromatin state or replication timing in determining the rate of mutation (likelihood ratio tests, all *P* > 0.15). Combining treatment groups, point mutations were associated with lower G/C content (*z* = –2.19, *P* = 0.028) and higher crossover recombination rates (*z* = 2.19, *P* = 0.028), with no significant effect of replication timing (*z* = –0.89, *P* = 0.37) or chromatin state (likelihood ratio test, *P* = 0.80) (Fig. 5). TE activity was associated with lower G/C content (*z* = –2.76, *P* = 5.75 × 10^−3^) and differed significantly among chromatin states (likelihood ratio test, *P* = 0.04), but did not vary with crossover recombination rate (*z* = –1.52, *P* = 0.13) or replication timing (*z* = –0.25, *P* = 0.80) (Fig. 6).

**Figure 5.**
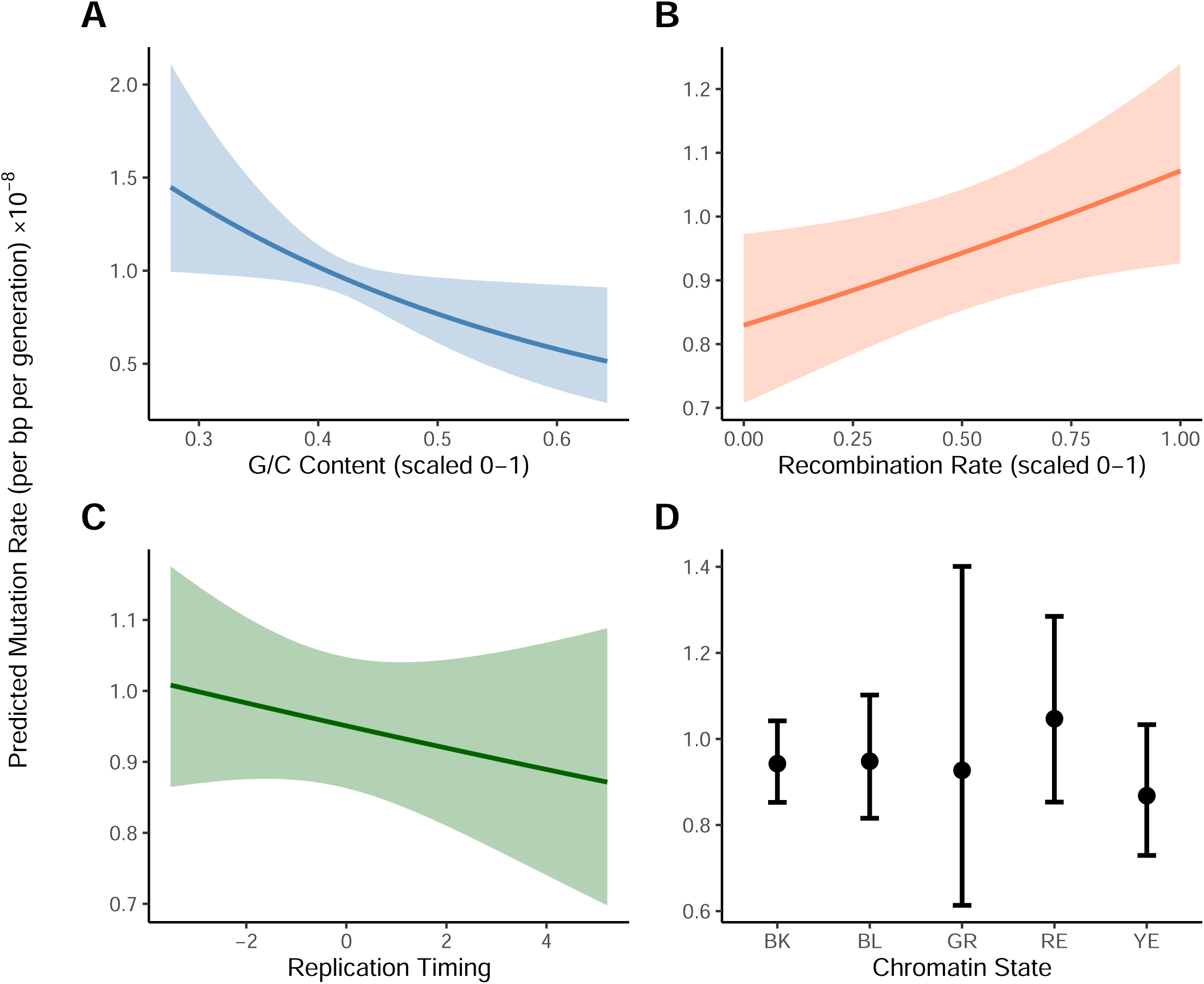
Relationships between point mutations and genomic context. Results are based on regression analysis of 7657 genomic windows, combining data across treatment groups, with 95% error bands. Point mutations were more likely to occur in regions of lower G/C content (A) and in regions of high crossover recombination (B). Point mutations were not significantly associated with replication timing (C) or chromatin state (D).

**Figure 6.**
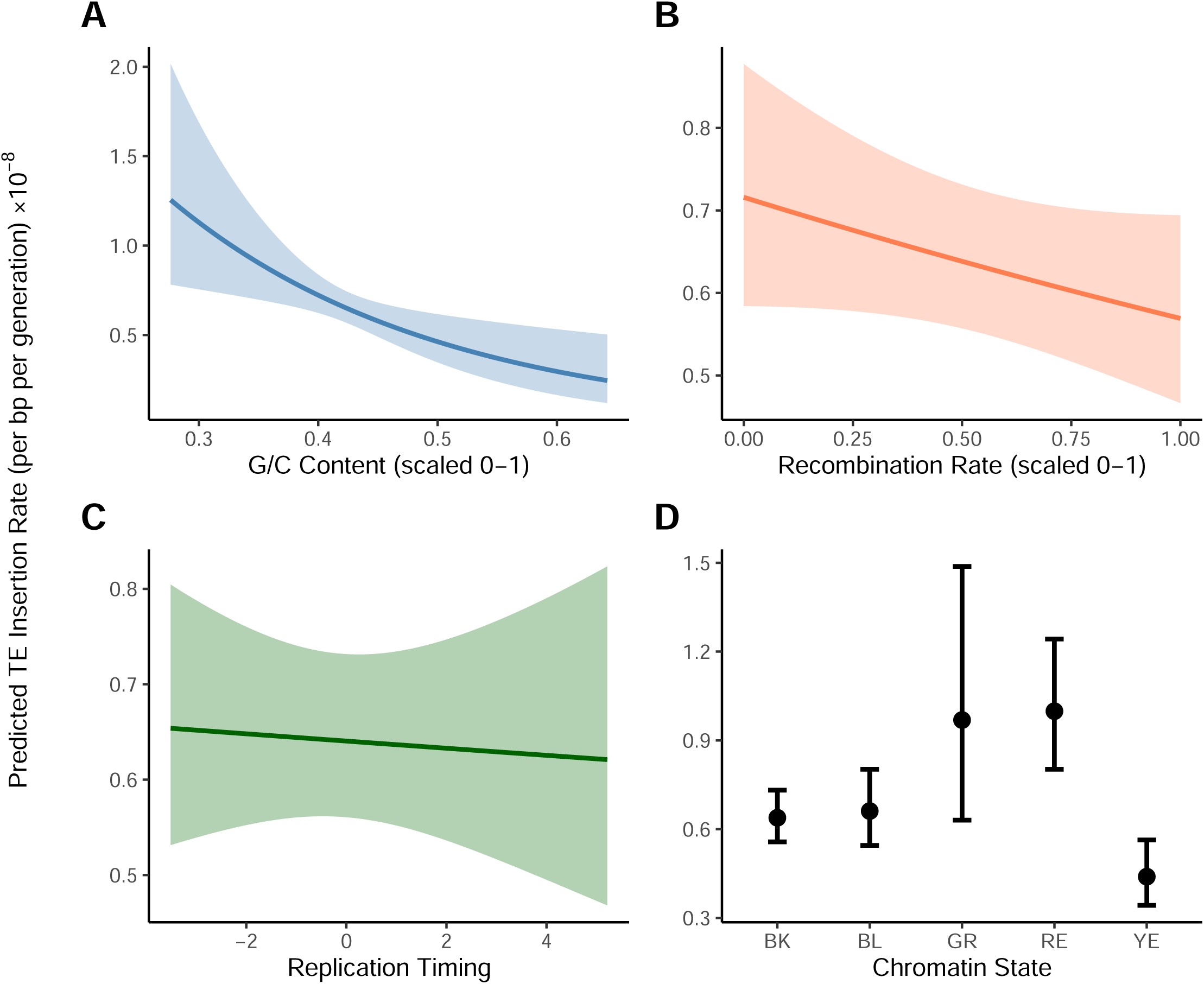
Relationships between transposable element activity and genomic context. Results are based on regression analysis of 7657 genomic windows, combining data across treatment groups, with 95% error bands. TE insertions were associated with lower G/C content (A) and chromatin state (D). TE insertions were not significantly associated with crossover recombination rate (B) or replication timing (C).

## Discussion

Eliminating endogenous meiotic DSBs appeared to have little effect on the overall rate of point mutations, but a substantial effect on the rate of transposition (Fig. 1). Our experimental design focused on genome-wide mutation patterns in the presence or absence of meiotic DSBs, rather than on mutations occurring at known or controlled sites of meiotic exchange. This approach permitted detection of ‘global’ effects–– transposition throughout the genome––but likely limited our power to detect mutation patterns associated with specific DSB locations. While not significant, we do find that the rate of single-nucleotide changes was greater in flies with endogenous DSBs, as expected if DSB repair is mutagenic. If we assume 24 meiotic DSBs per genome per generation (Carpenter, 1987; Mehrotra & McKim, 2006), and that each repair tract involves the synthesis of 5000 bp (Adams et al., 2003), our point estimates would be consistent with an increased mutation rate within such tracts of 6.7-fold.

We considered whether the presence of endogenous DSBs might disproportionately affect point mutation patterns in specific regions of the genome, particularly euchromatic regions where DSBs are more likely to occur, or areas of the genome showing higher levels of crossover recombination; we did not detect significant interactions with treatment for either of these factors. Instead, we found that point mutations were associated with regions of higher crossing over regardless of the presence of meiotic DSBs (Fig. 5), suggesting that some regions of the genome are more likely to experience both meiotic exchange and mutation, although the underlying mechanism for this is unclear. Such a pattern would contribute to the association between nucleotide diversity and crossover rate, independently of any mutagenic effect of DSBs. Importantly, we did not attempt to call point mutations in repetitive regions, which likely have distinct mutation patterns (Flynn et al., 2018; López-Cortegano et al., 2025; Nishant et al., 2009). An additional caveat to this analysis is that it incorporates data collected in separate experiments using cell lines and inbred strains that may or may not reflect the genomic features of the flies we studied.

An increased risk of mutation during the resolution of DSBs could be mitigated if the proteins that create DSBs also facilitate DNA repair, including the repair of exogenous, non-programmed DSBs. The suppression of endogenous DSBs in *mei-P22^−^* flies could limit the activity of DSB-dependent repair mechanisms such as the GATOR complex, potentially dampening responses to exogenous DNA damage (Wei et al., 2019), and counteracting any anti-mutagenic effect of eliminating endogenous DSBs.

In contrast to point mutations, we found a clear effect of our treatment on TE activity, with the rate of novel TE insertions increasing by 41% in *mei-P22^−^* flies (Fig. 1B). This increase was not restricted to a particular TE family (Fig. 2) or genomic context; TE events were associated with lower G/C content and certain chromatin states regardless of treatment (Fig. 6). One possible interpretation of this pattern is that *mei-P22^−^* flies were “stressed”, inducing TE activity, but only specific forms of stress appear to have such an effect in *Drosophila* (de Oliveira et al., 2021; Merenciano et al., 2025; Milyaeva et al., 2023; Mombach et al., 2022; Sharp & Agrawal, 2016). Instead, our results are consistent with previous observations linking TE activity with meiotic DSBs. Alleles disrupting the GATOR complex, which is involved in DSB repair, cause increased levels of TE expression (Wei et al., 2019), and alleles preventing proper synaptonemal complex formation result in fewer meiotic DSBs and increased TE insertion genome-wide (Miller, 2020). There is also a trend towards increased transposition in males, where endogenous DSBs are absent (Wang et al., 2023). TE repression may therefore be connected with the generation and repair of meiotic DSBs.

We found that TE insertions were more likely to occur on the X chromosome, in G/C poor regions, and in certain chromatin states (Fig. 6), matching previous findings (Adrion et al., 2017). However, the insertion bias towards certain chromatin states does not seem to be based simply on accessibility; insertion rates in the two types of euchromatin, RE and YE, both differed significantly from the highly repressed BK chromatin, but did so in opposite directions (Fig. 6). The continual improvement of bioinformatic tools for detecting TE movement will aid further investigation into complex insertion site preferences and distinguishing genomic patterns of TE mutation versus selection (Sultana et al., 2017; Zhang et al., 2020).

In conclusion, our results indicate that point mutations arising due to meiotic DSB repair represent a small fraction of the genome-wide mutational burden in *D. melanogaster*, and that spatial correlations between mutation and recombination may occur even in the absence of meiotic DSB repair. Our study also adds to the growing evidence that meiotic recombination in this organism is tightly associated with the suppression of TE activity.

## Data availability

Whole genome sequence data is available from the NCBI SRA under bioproject accession number PRJNA1346113. The R script used for analyses and its subsequent R workspace, our point mutation callset, TE callset, SV callset, the genomic windows used in analyses, and supplemental figures are available through Dryad DOI: 10.5061/dryad.b2rbnzstx.

## Acknowledgements

We would like to thank Denise Smith for performing the DNA extractions for this experiment. We would also like to thank Cindy Dong who assisted with general fly lab maintenance during the facilitation of this experiment.

## References

1. Adams, M. D., McVey, M., & Sekelsky, J. J. (2003). Drosophila BLM in Double-Strand Break Repair by Synthesis-Dependent Strand Annealing. Science, 299(5604), 265–267. 10.1126/science.1077198

2. Adrian, A. B., & Comeron, J. M. (2013). The Drosophila early ovarian transcriptome provides insight to the molecular causes of recombination rate variation across genomes. BMC Genomics, 14(1), 794. 10.1186/1471-2164-14-794

3. Adrian, A. B., Corchado, J. C., & Comeron, J. M. (2016). Predictive Models of Recombination Rate Variation across the Drosophila melanogaster Genome. Genome Biology and Evolution, 8(8), 2597–2612. 10.1093/gbe/evw181

4. Adrion, J. R., Song, M. J., Schrider, D. R., Hahn, M. W., & Schaack, S. (2017). Genome-Wide Estimates of Transposable Element Insertion and Deletion Rates in Drosophila Melanogaster. Genome Biology and Evolution, 9(5), 1329–1340. 10.1093/gbe/evx050

5. Andrews, S. (2010). FastQC: A Quality Control Tool for High Throughput Sequence Data [Computer software].

6. Baudat, F., Buard, J., Grey, C., Fledel-Alon, A., Ober, C., Przeworski, M., Coop, G., & de Massy, B. (2010). PRDM9 is a Major Determinant of Meiotic Recombination Hotspots in humans and mice. Science (New York, N.Y.), 327(5967), 836–840. 10.1126/science.1183439

7. Begun, D. J., & Aquadro, C. F. (1992). Levels of naturally occurring DNA polymorphism correlate with recombination rates in D. melanogaster. Nature, 356(6369), Article 6369. 10.1038/356519a0

8. Beyter, D., Ingimundardottir, H., Oddsson, A., Eggertsson, H. P., Bjornsson, E., Jonsson, H., Atlason, B. A., Kristmundsdottir, S., Mehringer, S., Hardarson, M. T., Gudjonsson, S. A., Magnusdottir, D. N., Jonasdottir, A., Jonasdottir, A., Kristjansson, R. P., Sverrisson, S. T., Holley, G., Palsson, G., Stefansson, O. A., … Stefansson, K. (2021). Long-read sequencing of 3,622 Icelanders provides insight into the role of structural variants in human diseases and other traits. Nature Genetics, 53(6), 779–786. 10.1038/s41588-021-00865-4

9. Biemont, C., Tsitrone, A., Vieira, C., & Hoogland, C. (1997). Transposable Element Distribution in Drosophila. Genetics, 147(4), 1997–1999. 10.1093/genetics/147.4.1997

10. Bolger, A. M., Lohse, M., & Usadel, B. (2014). Trimmomatic: A flexible trimmer for Illumina sequence data. Bioinformatics, 30(15), 2114–2120. 10.1093/bioinformatics/btu170

11. Broad Institute. (2019). Picard Toolkit [Computer software]. https://broadinstitute.github.io/picard/

12. Brooks, M. E., Kristensen, K., Benthem, K. J., van Magnusson, A., Berg, C. W., Nielsen, A., Skaug, H. J., Mächler, M., & Bolker, B. M. (2017). glmmTMB Balances Speed and Flexibility Among Packages for Zero-inflated Generalized Linear Mixed Modeling. The R Journal, 9(2), 378. 10.32614/rj-2017-066

13. Carpenter, A. T. C. (1987). Gene conversion, recombination nodules, and the initiation of meiotic synapsis. BioEssays, 6(5), 232–236. 10.1002/bies.950060510

14. Charlesworth, B., & Barton, N. H. (1996). Recombination load associated with selection for increased recombination. Genetics Research, 67(1), 27–41. 10.1017/S0016672300033450

15. Charlesworth, B., & Charlesworth, D. (1975). An experiment on recombination load in Drosophila melanogaster. Genetics Research, 25(3), 267–273. 10.1017/S001667230001569X

16. Chen, J., Basting, P. J., Han, S., Garfinkel, D. J., & Bergman, C. M. (2023). Reproducible evaluation of transposable element detectors with McClintock 2 guides accurate inference of Ty insertion patterns in yeast. Mobile DNA, 14, 8. 10.1186/s13100-023-00296-4

17. Chen, X., Schulz-Trieglaff, O., Shaw, R., Barnes, B., Schlesinger, F., Källberg, M., Cox, A. J., Kruglyak, S., & Saunders, C. T. (2016). Manta: Rapid detection of structural variants and indels for germline and cancer sequencing applications. Bioinformatics, 32(8), 1220–1222. 10.1093/bioinformatics/btv710

18. Comeron, J. M., Ratnappan, R., & Bailin, S. (2012). The Many Landscapes of Recombination in Drosophila melanogaster. PLOS Genetics, 8(10), e1002905. 10.1371/journal.pgen.1002905

19. Cutter, A. D., & Payseur, B. A. (2013). Genomic signatures of selection at linked sites: Unifying the disparity among species. Nature Reviews Genetics, 14(4), 262–274. 10.1038/nrg3425

20. Daigle, A., Whitehouse, L. S., Zhao, R., Emerson, J. J., & Schrider, D. R. (2025). Leveraging long-read assemblies and machine learning to enhance short-read transposable element detection and genotyping (p. 2025.02.11.637720). bioRxiv. 10.1101/2025.02.11.637720

21. de Oliveira, D. S., Rosa, M. T., Vieira, C., & Loreto, E. L. S. (2021). Oxidative and radiation stress induces transposable element transcription in Drosophila melanogaster. Journal of Evolutionary Biology, 34(4), 628–638. 10.1111/jeb.13762

22. Dolgin, E. S., & Charlesworth, B. (2008). The Effects of Recombination Rate on the Distribution and Abundance of Transposable Elements. Genetics, 178(4), 2169– 2177. 10.1534/genetics.107.082743

23. Dos Santos, G., Schroeder, A. J., Goodman, J. L., Strelets, V. B., Crosby, M. A., Thurmond, J., Emmert, D. B., Gelbart, W. M., & the FlyBase Consortium. (2015). FlyBase: Introduction of the Drosophila melanogaster Release 6 reference genome assembly and large-scale migration of genome annotations. Nucleic Acids Research, 43(D1), D690–D697. 10.1093/nar/gku1099

24. Dufourt, J., Dennis, C., Boivin, A., Gueguen, N., Théron, E., Goriaux, C., Pouchin, P., Ronsseray, S., Brasset, E., & Vaury, C. (2014). Spatio-temporal requirements for transposable element piRNA-mediated silencing during Drosophila oogenesis. Nucleic Acids Research, 42(4), 2512–2524. 10.1093/nar/gkt1184

25. Duina, A. A., Miller, M. E., & Keeney, J. B. (2014). Budding Yeast for Budding Geneticists: A Primer on the Saccharomyces cerevisiae Model System. Genetics, 197(1), 33–48. 10.1534/genetics.114.163188

26. Filion, G. J., van Bemmel, J. G., Braunschweig, U., Talhout, W., Kind, J., Ward, L. D., Brugman, W., de Castro, I. J., Kerkhoven, R. M., Bussemaker, H. J., & van Steensel, B. (2010). Systematic protein location mapping reveals five principal chromatin types in Drosophila cells. Cell, 143(2), 212–224. 10.1016/j.cell.2010.09.009

27. Flynn, J. M., Lower, S. E., Barbash, D. A., & Clark, A. G. (2018). Rates and Patterns of Mutation in Tandem Repetitive DNA in Six Independent Lineages of Chlamydomonas reinhardtii. Genome Biology and Evolution, 10(7), 1673–1686. 10.1093/gbe/evy123

28. Halldorsson, B. V., Palsson, G., Stefansson, O. A., Jonsson, H., Hardarson, M. T., Eggertsson, H. P., Gunnarsson, B., Oddsson, A., Halldorsson, G. H., Zink, F., Gudjonsson, S. A., Frigge, M. L., Thorleifsson, G., Sigurdsson, A., Stacey, S. N., Sulem, P., Masson, G., Helgason, A., Gudbjartsson, D. F., … Stefansson, K. (2019). Characterizing mutagenic effects of recombination through a sequence-level genetic map. Science, 363(6425), eaau1043. 10.1126/science.aau1043

29. Hardin, J. W., & Hilbe, J. M. (2007). Generalized Linear Models and Extensions, Second Edition. Stata Press.

30. Hinch, R., Donnelly, P., & Hinch, A. G. (2023). Meiotic DNA breaks drive multifaceted mutagenesis in the human germ line. Science (New York, N.Y.), 382(6674), eadh2531. 10.1126/science.adh2531

31. Huang, Y., Gao, Z. Y., Ly, K., Lin, L., Lambooij, J.-P., King, E. G., Janssen, A., Wei, K. H.-C., & Lee, Y. C. G. (2025). Polymorphic transposable elements contribute to variation in recombination landscapes. Proceedings of the National Academy of Sciences, 122(12), e2427312122. 10.1073/pnas.2427312122

32. Iida, T., & Lilly, M. A. (2004). Missing oocyte encodes a highly conserved nuclear protein required for the maintenance of the meiotic cycle and oocyte identity in Drosophila. Development, 131(5), 1029–1039. 10.1242/dev.01001

33. Jeffreys, A. J., Kauppi, L., & Neumann, R. (2001). Intensely punctate meiotic recombination in the class II region of the major histocompatibility complex. Nature Genetics, 29(2), 217–222. 10.1038/ng1001-217

34. Katju, V., & Bergthorsson, U. (2019). Old Trade, New Tricks: Insights into the Spontaneous Mutation Process from the Partnering of Classical Mutation Accumulation Experiments with High-Throughput Genomic Approaches. Genome Biology and Evolution, 11(1), 136–165. 10.1093/gbe/evy252

35. Keightley, P. D., & Otto, S. P. (2006). Interference among deleterious mutations favours sex and recombination in finite populations. Nature, 443(7107), 89–92. 10.1038/nature05049

36. Kelleher, E. S., Barbash, D. A., & Blumenstiel, J. P. (2020). Taming the Turmoil Within: New Insights on the Containment of Transposable Elements. Trends in Genetics: TIG, 36(7), 474–489. 10.1016/j.tig.2020.04.007

37. Kent, T. V., Uzunović, J., & Wright, S. I. (2017). Coevolution between transposable elements and recombination. Philosophical Transactions of the Royal Society B: Biological Sciences, 372(1736), 20160458. 10.1098/rstb.2016.0458

38. Koehler, K. E., Hawley, R. S., Sherman, S., & Hassold, T. (1996). Recombination and nondisjunction in humans and flies. Human Molecular Genetics, 5(Supplement_1), 1495–1504. 10.1093/hmg/5.Supplement_1.1495

39. Kofler, R., Betancourt, A. J., & Schlötterer, C. (2012). Sequencing of Pooled DNA Samples (Pool-Seq) Uncovers Complex Dynamics of Transposable Element Insertions in Drosophila melanogaster. PLOS Genetics, 8(1), e1002487. 10.1371/journal.pgen.1002487

40. Lake, C. M., Nielsen, R. J., Bonner, A. M., Eche, S., White-Brown, S., McKim, K. S., & Hawley, R. S. (2019). Narya, a RING finger domain-containing protein, is required for meiotic DNA double-strand break formation and crossover maturation in Drosophila melanogaster. PLoS Genetics, 15(1), e1007886. 10.1371/journal.pgen.1007886

41. Lake, C. M., Nielsen, R. J., Guo, F., Unruh, J. R., Slaughter, B. D., & Hawley, R. S. (2015). Vilya, a component of the recombination nodule, is required for meiotic double-strand break formation in Drosophila. eLife, 4, e08287. 10.7554/eLife.08287

42. Lake, C. M., Nielsen, R. J., & Hawley, R. S. (2011). The Drosophila Zinc Finger Protein Trade Embargo Is Required for Double Strand Break Formation in Meiosis. PLOS Genetics, 7(2), e1002005. 10.1371/journal.pgen.1002005

43. Lee, L., & Rosin, L. F. (2024). Uncharted territories: Solving the mysteries of male meiosis in flies. PLOS Genetics, 20(3), e1011185. 10.1371/journal.pgen.1011185

44. Li, H., Handsaker, B., Wysoker, A., Fennell, T., Ruan, J., Homer, N., Marth, G., Abecasis, G., Durbin, R., & 1000 Genome Project Data Processing Subgroup. (2009). The Sequence Alignment/Map format and SAMtools. Bioinformatics, 25(16), 2078–2079. 10.1093/bioinformatics/btp352

45. Liu, H., Jang, J. K., Kato, N., & McKim, K. S. (2002). Mei-P22 Encodes a Chromosome-Associated Protein Required for the Initiation of Meiotic Recombination in Drosophila melanogaster. Genetics, 162(1), 245–258. 10.1093/genetics/162.1.245

46. Liu, H., Jia, Y., Sun, X., Tian, D., Hurst, L. D., & Yang, S. (2017). Direct Determination of the Mutation Rate in the Bumblebee Reveals Evidence for Weak Recombination-Associated Mutation and an Approximate Rate Constancy in Insects. Molecular Biology and Evolution, 34(1), 119–130. 10.1093/molbev/msw226

47. Longhese, M. P., Bonetti, D., Guerini, I., Manfrini, N., & Clerici, M. (2009). DNA double-strand breaks in meiosis: Checking their formation, processing and repair. DNA Repair, 8(9), 1127–1138. 10.1016/j.dnarep.2009.04.005

48. López-Cortegano, E., Chebib, J., Jonas, A., Vock, A., Künzel, S., Keightley, P. D., & Tautz, D. (2025). The rate and spectrum of new mutations in mice inferred by long-read sequencing. Genome Research, 35(1), 43–54. 10.1101/gr.279982.124

49. Magni, G. E. (1963). The origin of spontaneous mutations during meiosis. Proceedings of the National Academy of Sciences, 50(5), 975–980. 10.1073/pnas.50.5.975

50. McKim, K. S., Green-Marroquin, B. L., Sekelsky, J. J., Chin, G., Steinberg, C., Khodosh, R., & Hawley, R. S. (1998). Meiotic Synapsis in the Absence of Recombination. Science, 279(5352), 876–878. 10.1126/science.279.5352.876

51. McLaren, W., Gil, L., Hunt, S. E., Riat, H. S., Ritchie, G. R. S., Thormann, A., Flicek, P., & Cunningham, F. (2016). The Ensembl Variant Effect Predictor. Genome Biology, 17(1), 122. 10.1186/s13059-016-0974-4

52. Mehrotra, S., & McKim, K. S. (2006). Temporal Analysis of Meiotic DNA Double-Strand Break Formation and Repair in Drosophila Females. PLOS Genetics, 2(11), e200. 10.1371/journal.pgen.0020200

53. Merenciano, M., Oliveira, D. S., Salces-Ortiz, J., Rebollo, R., Manfré, B., Menezes, B., Krasovec, G., Simonet, C., Janillon, S., Burlet, N., Carareto, C. M. A., Vieira, C., & Fablet, M. (2025). Gene and transposable element expression in response to stress in temperate and tropical populations of Drosophila. Mobile DNA, 16(1), 35. 10.1186/s13100-025-00372-x

54. Miller, D. E. (2020). Synaptonemal Complex-Deficient Drosophila melanogaster Females Exhibit Rare DSB Repair Events, Recurrent Copy-Number Variation, and an Increased Rate of de Novo Transposable Element Movement. G3 Genes|Genomes|Genetics, 10(2), 525–537. 10.1534/g3.119.400853

55. Miller, D. E., Cook, K. R., Yeganeh Kazemi, N., Smith, C. B., Cockrell, A. J., Hawley, R. S., & Bergman, C. M. (2016). Rare recombination events generate sequence diversity among balancer chromosomes in Drosophila melanogaster. Proceedings of the National Academy of Sciences, 113(10), E1352–E1361. 10.1073/pnas.1601232113

56. Milyaeva, P. A., Kukushkina, I. V., Kim, A. I., & Nefedova, L. N. (2023). Stress Induced Activation of LTR Retrotransposons in the Drosophila melanogaster Genome. Life, 13(12), 2272. 10.3390/life13122272

57. Mombach, D. M., da Fontoura Gomes, T. M. F., & Loreto, E. L. S. (2022). Stress does not induce a general transcription of transposable elements in Drosophila. Molecular Biology Reports, 49(9), 9033–9040. 10.1007/s11033-022-07839-7

58. Nelson, M. G., Linheiro, R. S., & Bergman, C. M. (2017). McClintock: An Integrated Pipeline for Detecting Transposable Element Insertions in Whole-Genome Shotgun Sequencing Data. G3: Genes|Genomes|Genetics, 7(8), 2763–2778. 10.1534/g3.117.043893

59. Nishant, K. T., Singh, N. D., & Alani, E. (2009). Genomic mutation rates: What high-throughput methods can tell us. BioEssays: News and Reviews in Molecular, Cellular and Developmental Biology, 31(9), 912–920. 10.1002/bies.200900017

60. Petronczki, M., Siomos, M. F., & Nasmyth, K. (2003). Un Ménage à Quatre: The Molecular Biology of Chromosome Segregation in Meiosis. Cell, 112(4), 423–440. 10.1016/S0092-8674(03)00083-7

61. Petrov, D. A., Fiston-Lavier, A.-S., Lipatov, M., Lenkov, K., & González, J. (2011). Population Genomics of Transposable Elements in Drosophila melanogaster. Molecular Biology and Evolution, 28(5), 1633–1644. 10.1093/molbev/msq337

62. Pratto, F., Brick, K., Khil, P., Smagulova, F., Petukhova, G. V., & Camerini-Otero, R. D. (2014). DNA recombination. Recombination initiation maps of individual human genomes. *Science (New York*, N.Y.), 346(6211), 1256442. 10.1126/science.1256442

63. R Core Team. (2023). R: A Language and Environment for Statistical Computing [Computer software]. R Foundation for Statistical Computing. https://www.R-project.org/

64. Rattray, A., Santoyo, G., Shafer, B., & Strathern, J. N. (2015). Elevated Mutation Rate during Meiosis in Saccharomyces cerevisiae. PLOS Genetics, 11(1), e1004910. 10.1371/journal.pgen.1004910

65. Ray, D., & Ye, P. (2013). Characterization of the Metabolic Requirements in Yeast Meiosis. PLOS ONE, 8(5), e63707. 10.1371/journal.pone.0063707

66. Rech, G. E., Radío, S., Guirao-Rico, S., Aguilera, L., Horvath, V., Green, L., Lindstadt, H., Jamilloux, V., Quesneville, H., & González, J. (2022). Population-scale long-read sequencing uncovers transposable elements associated with gene expression variation and adaptive signatures in Drosophila. Nature Communications, 13, 1948. 10.1038/s41467-022-29518-8

67. Rodgers, K., & McVey, M. (2016). Error-Prone Repair of DNA Double-Strand Breaks. Journal of Cellular Physiology, 231(1), 15–24. 10.1002/jcp.25053

68. Schwaiger, M., Stadler, M. B., Bell, O., Kohler, H., Oakeley, E. J., & Schübeler, D. (2009). Chromatin state marks cell-type– and gender-specific replication of the Drosophila genome. Genes & Development, 23(5), 589–601. 10.1101/gad.511809

69. Sekelsky, J. J., McKim, K. S., Messina, L., French, R. L., Hurley, W. D., Arbel, T., Chin, G. M., Deneen, B., Force, S. J., Hari, K. L., Jang, J. K., Laurençon, A. C., Madden, L. D., Matthies, H. J., Milliken, D. B., Page, S. L., Ring, A. D., Wayson, S. M., Zimmerman, C. C., & Hawley, R. S. (1999). Identification of Novel Drosophila Meiotic Genes Recovered in a P-Element Screen. Genetics, 152(2), 529–542. 10.1093/genetics/152.2.529

70. Sharp, N. P., & Agrawal, A. F. (2016). Low Genetic Quality Alters Key Dimensions of the Mutational Spectrum. PLOS Biology, 14(3), e1002419. 10.1371/journal.pbio.1002419

71. Smit, A., Hubley, R., & Green, P. (2013). RepeatMasker Open-4.0 [Computer software]. http://www.repeatmasker.org

72. Smukowski, C. S., & Noor, M. a. F. (2011). Recombination rate variation in closely related species. Heredity, 107(6), 496–508. 10.1038/hdy.2011.44

73. Strathern, J. N., Shafer, B. K., & McGill, C. B. (1995). DNA synthesis errors associated with double-strand-break repair. Genetics, 140(3), 965–972. 10.1093/genetics/140.3.965

74. Sultana, T., Zamborlini, A., Cristofari, G., & Lesage, P. (2017). Integration site selection by retroviruses and transposable elements in eukaryotes. Nature Reviews Genetics, 18(5), 292–308. 10.1038/nrg.2017.7

75. Van der Auwera, G., & O’Connor, B. (2020). Genomics in the Cloud (1st edition). O’Reilly Media, Incorporated.

76. Vasimuddin, Md., Misra, S., Li, H., & Aluru, S. (2019). Efficient Architecture-Aware Acceleration of BWA-MEM for Multicore Systems. 2019 IEEE International Parallel and Distributed Processing Symposium (IPDPS), 314–324. 10.1109/IPDPS.2019.00041

77. Wang, Y., McNeil, P., Abdulazeez, R., Pascual, M., Johnston, S. E., Keightley, P. D., & Obbard, D. J. (2023). Variation in mutation, recombination, and transposition rates in Drosophila melanogaster and Drosophila simulans. Genome Research, 33(4), 587–598. 10.1101/gr.277383.122

78. Wei, Y., Bettedi, L., Ting, C.-Y., Kim, K., Zhang, Y., Cai, J., & Lilly, M. A. (2019). The GATOR complex regulates an essential response to meiotic double-stranded breaks in Drosophila. eLife, 8, e42149. 10.7554/eLife.42149

79. Wylie, A., Jones, A. E., D’Brot, A., Lu, W.-J., Kurtz, P., Moran, J. V., Rakheja, D., Chen, K. S., Hammer, R. E., Comerford, S. A., Amatruda, J. F., & Abrams, J. M. (2016). P53 genes function to restrain mobile elements. Genes & Development, 30(1), 64–77. 10.1101/gad.266098.115

80. Yang, S., Wang, L., Huang, J., Zhang, X., Yuan, Y., Chen, J.-Q., Hurst, L. D., & Tian, D. (2015). Parent–progeny sequencing indicates higher mutation rates in heterozygotes. Nature, 523(7561), 463–467. 10.1038/nature14649

81. Yu, T., Huang, X., Dou, S., Tang, X., Luo, S., Theurkauf, W. E., Lu, J., & Weng, Z. (2021). A benchmark and an algorithm for detecting germline transposon insertions and measuring de novo transposon insertion frequencies. Nucleic Acids Research, 49(8), e44. 10.1093/nar/gkab010

82. Zhang, X., Zhao, M., McCarty, D. R., & Lisch, D. (2020). Transposable elements employ distinct integration strategies with respect to transcriptional landscapes in eukaryotic genomes. Nucleic Acids Research, 48(12), 6685–6698. 10.1093/nar/gkaa370

